# Assessing cross-contamination in spike-sorted electrophysiology data

**DOI:** 10.1101/2023.12.21.572882

**Authors:** Jack P. Vincent, Michael N. Economo

## Abstract

Recent advances in extracellular electrophysiology now facilitate the recording of spikes from hundreds or thousands of neurons simultaneously. This has necessitated both the development of new computational methods for spike sorting and better methods to determine spike sorting accuracy. One longstanding method of assessing the false discovery rate (FDR) of spike sorting – the rate at which spikes are misassigned to the wrong cluster – has been the rate of inter-spike-interval (ISI) violations. Despite their near ubiquitous usage in spike sorting, our understanding of how exactly ISI violations relate to FDR, as well as best practices for using ISI violations as a quality metric, remain limited. Here, we describe an analytical solution that can be used to predict FDR from ISI violation rate. We test this model in silico through Monte Carlo simulation, and apply it to publicly available spike-sorted electrophysiology datasets. We find that the relationship between ISI violation rate and FDR is highly nonlinear, with additional dependencies on firing rate, the correlation in activity between neurons, and contaminant neuron count. Predicted median FDRs in public datasets were found to range from 3.1% to 50.0%. We find that stochasticity in the occurrence of ISI violations as well as uncertainty in cluster-specific parameters make it difficult to predict FDR for single clusters with high confidence, but that FDR can be estimated accurately across a population of clusters. Our findings will help the growing community of researchers using extracellular electrophysiology assess spike sorting accuracy in a principled manner.

**SIGNIFICANCE STATEMENT:** High-density silicon probes are widely used to record the activity of large populations of neurons while animals are engaged in complex behavior. In this approach, each electrode records spikes from many neurons, and “spike sorting” algorithms are used to group the spikes originating from each neuron together. This process is error-prone, however, and so the ability to assess spike sorting accuracy is essential for properly interpreting neural activity. The rate of inter-spike-interval (ISI) violations is commonly used to assess spike sorting accuracy, but the relationship between ISI violation rate and sorting accuracy is complex and poorly understood. Here, we describe this relationship in detail and provide guidelines for how to properly use ISI violation rate to assess spike sorting accuracy.

## INTRODUCTION

Extracellular electrophysiology has become an increasingly popular method for studying neuronal activity at the population level. Silicon probes containing dozens or hundreds of densely packed electrode sites can be used to observe neuronal action potentials in many neurons simultaneously. This multiplexed signal acquisition, however, often necessitates the assignment of observed action potentials to individual neurons - “spike sorting” - as a critical first step prior to many analyses (Rey et al., 2015; Todorova et al., 2014). Approaches for spike sorting vary, though all generally involve comparisons of observed action potential waveforms within and across electrodes, and then grouping similar spikes – thought to be produced by the same neuron – together, forming clusters (Gibson et al., 2012; Quiroga & Panzeri, 2013). Ideally, each cluster is composed principally of true positive (TP) spikes from a single neuron making it a “well isolated” cluster. Spikes from this single neuron can be erroneously excluded from the cluster, resulting in false negatives (FNs). The primary focus of this work, however, is false positives (FPs), wherein spikes are misassigned to a cluster whose activity is meant to correspond to a different neuron. Clusters with substantial “contamination” by FPs thus represent the combined activity of multiple neurons.

False positives are a persistent problem in spike sorting that results from frequently unavoidable similarities in action potential waveforms, occurrence of spikes at overlapping times, nonstationarity in waveform shape, and recording noise. Contamination can distort the activity of a cluster, potentially leading to incorrect conclusions about how single neurons encode information. For instance, a population of neurons recorded during a task with two cues may contain neurons responsive to just one cue or the other. However, poor sorting may lead to cluster cross-contamination between these two phenotypes, giving the impression that neurons in this region respond to both cues. The prevalence of FPs in a single cluster, recording session, or dataset can be described by the false discovery rate (FDR), a value that ranges between 0 and 1 and reports the proportion of sorted spikes that have been misassigned. While FNs can also be a concern, as they can decrease recorded spike rates and thus reduce statistical power in subsequent analyses (Hill et al., 2011), FNs should not generally alter the overall patterns of activity associated with individual clusters and are thus of less concern than FPs.

Algorithmic approaches to spike sorting for high-density silicon probe electrophysiology have seen concentrated development over the last decade, with researchers now selecting from a number of competing options for high-throughput automated spike sorting (Bestel et al., 2012; Buccino et al., 2020; Chaure et al., 2018; Chung et al., 2017; Jun et al., 2017; J. H. Lee et al., 2017; Pachitariu et al., 2016; Saif-ur-Rehman et al., 2021; Toosi et al., 2021), yet concomitant techniques for *post hoc* analysis of spike-sorted data (Barnett et al., 2016; Hill et al., 2011; Magland et al., 2020; Neymotin et al., 2011; Pouzat et al., 2002) have been given comparatively less attention. Sorting quality metrics generally fall into two categories: assessment of cluster overlap using dimensionally reduced representations of sorted spike waveforms, e.g. L-ratio (Schmitzer-Torbert et al., 2005), isolation distance (Harris et al., 2001), D-prime (Hill et al., 2011), and silhouette score, or empirical measures known to be related to cluster isolation and sorting difficulty, e.g. signal-to-noise ratio, presence ratio, firing range, and inter-spike interval (ISI) violations. Among all these metrics, ISI violations are unique in that they are tightly linked to the occurrence of FPs. Biophysical limitations prevent neurons from producing consecutive spikes within their absolute refractory period, meaning the presence of action potentials spaced by less than the absolute refractory period, an ISI violation, is always the result of at least one FP.

ISI violations are typically reported as a fraction of the total number of spikes assigned to a cluster. The ISI violation rate (ISI_v_) – the number of ISI violations divided by the total number of spikes assigned to a cluster – is often interpreted subjectively with only a general understanding that a lower ISI_v_ is associated with a lower FDR. Often, although not always, it is appreciated that most FP spikes do not produce ISI violations and so ISI_v_ << FDR. Previous work has estimated the relationship between ISI_v_ and FDR (Hill et al., 2011; Llobet et al., 2022) using simplifying assumptions, but the accuracy and limitations of predicting cluster FDR on the basis of ISI violations under realistic experimental conditions have not been assessed.

Here, we developed a comprehensive model explaining the relationship between ISI_v_ and FDR following spike sorting with respect to cluster contamination, neuronal firing rate, the temporal relationships between neurons, and the number of neurons contributing FPs. We benchmark the accuracy of this model *in silico* through Monte Carlo simulation and explore limitations in the accuracy of FDR estimation imposed by *in vivo* recording conditions. Finally, we apply this model to publicly available spike-sorted electrophysiology data to provide an estimate of FDRs based on ISI violations in the literature to provide researchers intuition about the expected range of FDR associated with silicon probe electrophysiology.

## MATERIALS AND METHODS

### Monte-Carlo simulation of neural spike trains

Stochastic neural spike trains were simulated using Elephant (Electrophysiology Analysis Toolkit) (Yegenoglu et al., 2018). Specifically, neurons were modeled as either homogeneous or inhomogeneous Poisson processes (van Vreeswijk, 2010) using either the StationaryPoissonProcess or the NonStationaryPoissonProcess functions of the spike_train_generation module. Custom Python scripts were used for subsequent simulation and analysis. A refractory period of 2.5 ms was assumed for all simulations as well as all calculations in *Table 1*. Almost all datasets examined were recorded from mouse cortical neurons, however for data collected from different cell types or animal models, this parameter would need to be adjusted accordingly. For generation of inhomogeneous spike trains, peristimulus time histograms (PSTHs) derived from several of the publicly available electrophysiology datasets examined in *Table 1* were used. Simulated recording durations varied depending on the desired level of certainty in ISI_v_ and need to emulate realistic recording conditions. These times included ∼28 hours (**Fig. 3A-B**), ∼17 hours (**Fig. 3C**), 12 hours (**Fig. 3D**), 30 min (**Fig. 2B**), and 10 min (**Fig. 2A**; **Fig. 2C**; **Fig.4B-C**)

**Table 1.** Median and mean FDR of publicly available spike-sorted electrophysiology datasets.

| Authors | Median FDR<br>(± s.e.) | Mean FDR<br>(± s.e.) | # Clusters | $\tau_c$<br>(ms) | Probe | Sorter |
| --- | --- | --- | --- | --- | --- | --- |
| Xu et al.<br>*Nature* (2022) | 3.1% (0.5) | 14.3% (0.4) | 3,046 | 0 | H2/H3<br>*Cam. Neuro.* | Kilosort |
| Economo et al.<br>*Nature* (2018) | 3.1% (0.6) | 12.5% (0.5) | 1,988 | 0.25 | H2<br>*Cam. Neuro.* | JRClust |
| Steinmetz et al.<br>*Nature* (2019)* | 4.0% (0.2) | 25.1% (0.2) | 33,997 | 0 | Neuropixels 1.0 | Kilosort |
| Gao et al.<br>*Nature* (2018) | 4.4% (0.7) | 18.7% (0.6) | 1,923 | 0.5 | H2/A4<br>*Cam. Neuro.* | UltraMegaSort2000 |
| Li et al.<br>*Nature* (2016) | 4.5% (0.8) | 19.7% (0.7) | 1,543 | 0.85 | A4<br>*NeuroNexus* | UltraMegaSort2000 |
| Inagaki et al.<br>*Cell* (2022) | 6.2% (0.3) | 17.7% (0.3) | 7,968 | 0.25 | HH-2/Neuropixels 2.0<br>*Janelia/N.A.* | JRClust/Kilosort |
| Guo et al.<br>*Nature* (2017) | 6.5% (0.7) | 18.8% (0.6) | 1,936 | 0.85 | A4/A2<br>*NeuroNexus* | UltraMegaSort2000 |
| Finkelstein et al.<br>*Nat. Neuro.* (2021)* | 10.5% (0.5) | 20.1% (0.4) | 3,385 | 0.25 | H2<br>*Cam. Neuro.* | JRClust |
| Sylwestrak et al.<br>*Cell* (2022) | 12.8% (0.4) | 29.1% (0.3) | 10,548 | 0.25 | Neuropixels 2.0 | Kilosort |
| Stringer et al.<br>*Science* (2019)<sup>†</sup> | 21.9% (0.5) | 33.1% (0.4) | 6,446 | 0 | Neuropixels 1.0 | Kilosort |
| Chinta & Pluta<br>*Nat. Comm.* (2023) | 28.2% (1.3) | 36.2% (1.0) | 991 | 0 | NeuroNexus | Kilosort |
| Juavinett et al.<br>*eLife* (2019)*<sup>†</sup> | 50.0% (0.8) | 44.3% (0.7) | 2,018 | 0 | Neuropixels 1.0 | Kilosort |

\* Clusters labeled “multi” or “MUA” excluded
<sup>†</sup> Calculated using homogeneous equation (Eq. 9)

**Figure 1.**
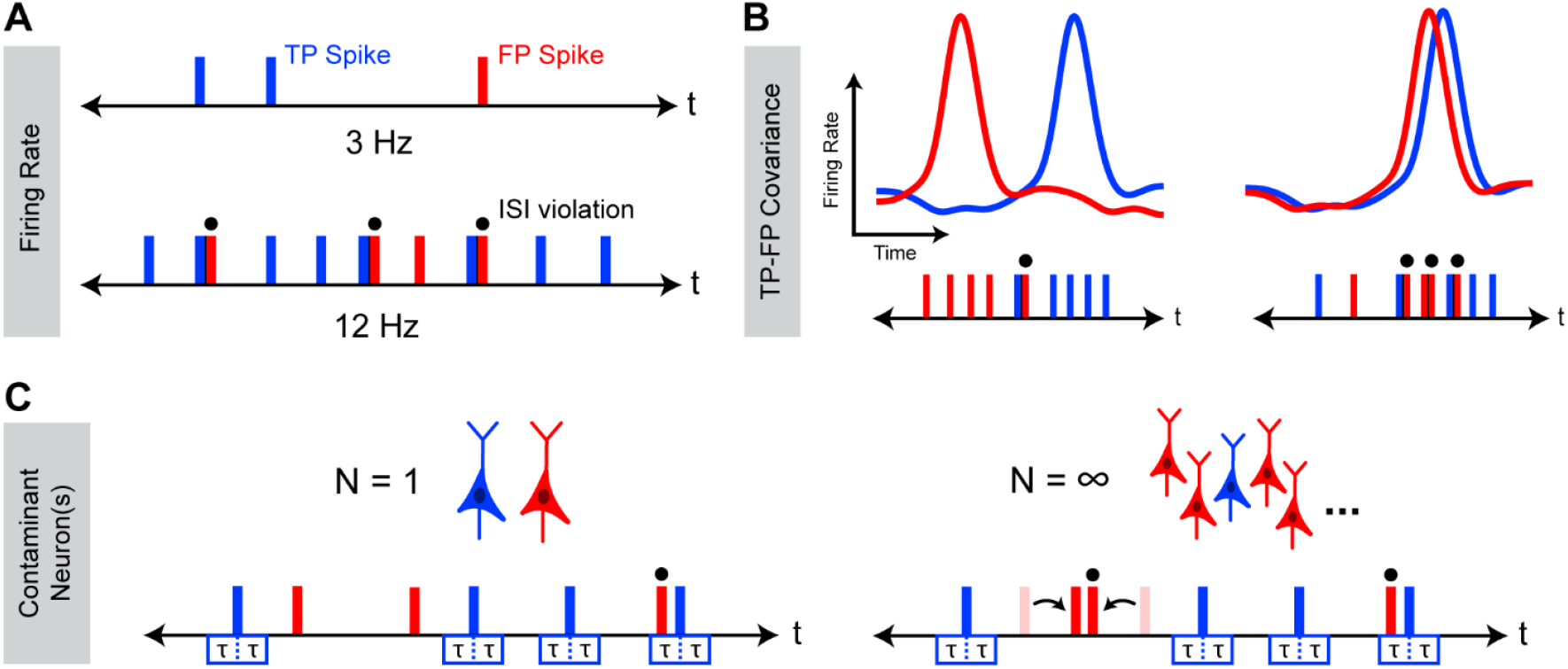
Factors affecting the relationship between ISI_v_ and FDR. Schematic representation of occurrence of ISI violations for a cluster with varying firing rates (**A**) TP-FP covariance (**B**), and numbers of contaminant neuron(s) (**C**). In all cases, underlying FDR between the two cases is the same, while observed ISI_v_ varies as a consequence of changes in these characteristics of neuronal activity. Blue corresponds to true positive (TP) spikes, and red corresponds to false positive (FP) spikes. Overhead dots indicate observed ISI violations. τ represents the neuronal refractory period.

**Figure 2.**
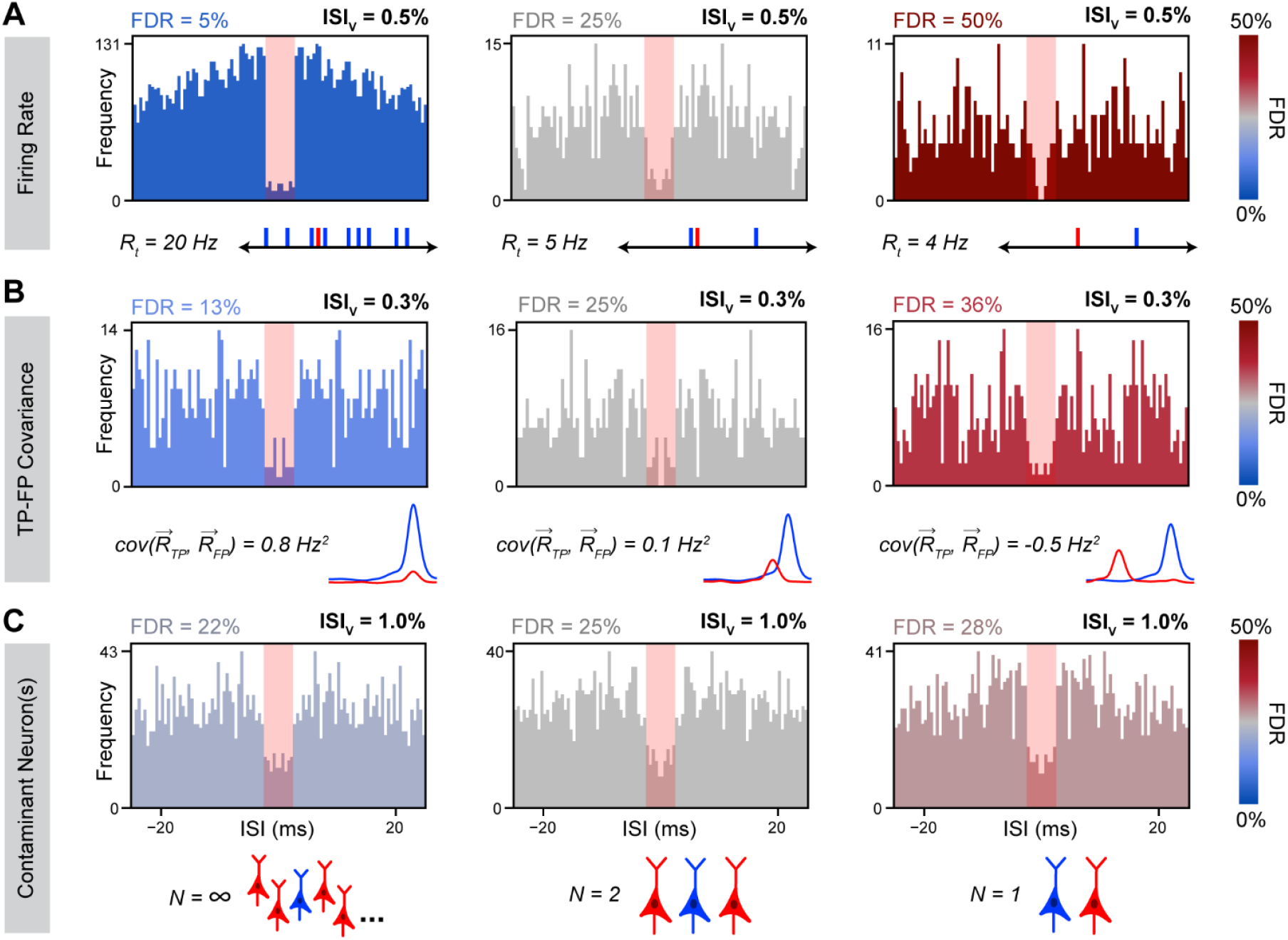
Simulated ISI distributions with variable underlying neuronal characteristics. Representative ISI histograms with all factors kept constant except total cluster firing rate (**A**), TP-FP covariance (**B**) and contaminant neuron count (**C**). Simulated firing rates were as indicated (*A*), 2 Hz (*B*), and 10 Hz (*C*). Homogeneous firing assumed for the top and bottom rows. Blue corresponds to true positive (TP) spikes, and red corresponds to false positive (FP) spikes. FP firing rate traces (*B*) are schematics only.

**Figure 3.**
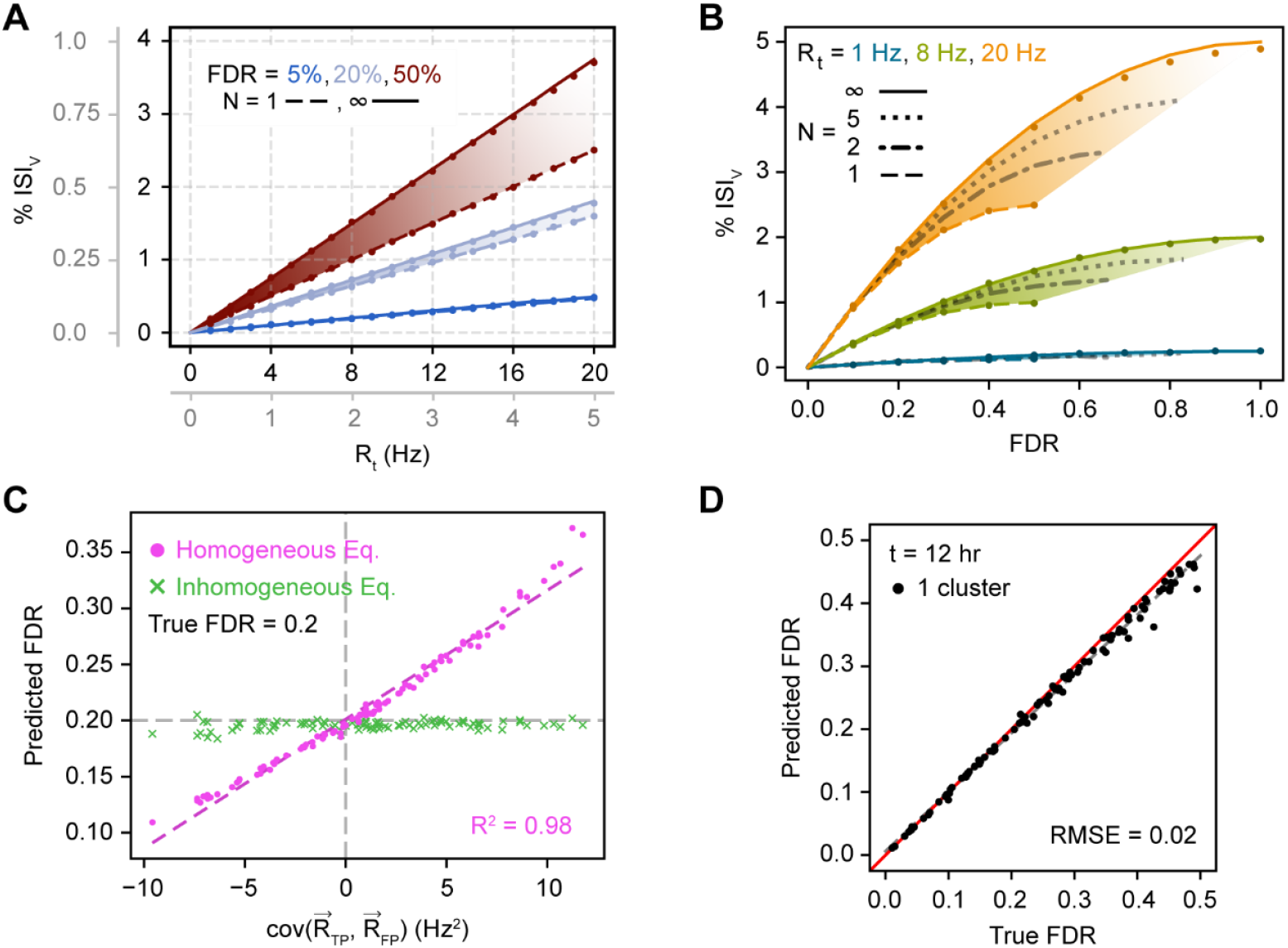
Relationship between single cluster FDR and observed ISI_v_. (**A**) Dependence of ISI_v_ on total firing rate given varying FDRs and contaminant neuron counts. Lines correspond to analytical predictions; dots correspond to simulation results. Plotted data applies to both primary and gray axes. (**B**) Dependence of ISI_v_ on FDR given varying firing rates and contaminant neuron counts. Conventions as in (*A*). (**C**) Prediction of FDR from observed ISI_v_ with temporally inhomogeneous firing rates using either the homogeneous model (*Eq. 9*) or the inhomogeneous model (*Eq. 10*). (**D**) Prediction of FDR from observed ISI_v_ across a range of physiologically relevant underlying neuronal characteristics (see *Materials and Methods*) for 100 total simulated clusters. Red line is the unity line, or perfect concurrence between predicted and true FDR; dashed gray line is the line of best fit. RMSE calculated with respect to the unity line.

### Monte Carlo simulation of cluster populations

In some simulations, all parameters were varied simultaneously to produce populations of clusters with a wide range of physiologically feasible parametric combinations (**Fig. 3D**; **Fig. 4B-C**). For each cluster, FDR was sampled from bounded Cauchy distributions, as these were found to most accurately replicate ISI_v_ distributions found in electrophysiological data when simulated. Different population median and mean FDRs were obtained by adjusting the location and scale of the Cauchy distribution. R_t_ was randomly selected from bounded uniform distributions: [4, 20] (**Fig. 3D**), [1, 10] (**Fig. 4B**), and [4, 16] (**Fig. 4C**). Possible values of N were randomly chosen from among the following values: [1, 2, 5, ∞]. This was found to be the most balanced way to sample N, as the effect of contaminant neuron count on ISI violation occurrence increases logarithmically. Lastly TP-FP covariance was varied by randomly selecting peristimulus time histograms (PSTHs) from electrophysiology datasets (**Table 1**) to serve as 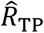 and 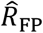. These PSTHs were then scaled appropriately and combined to reach the desired R_t_ and FDR. TP-FP covariance using this method varied from -49.2 to 113.7 Hz^2^.

**Figure 4.**
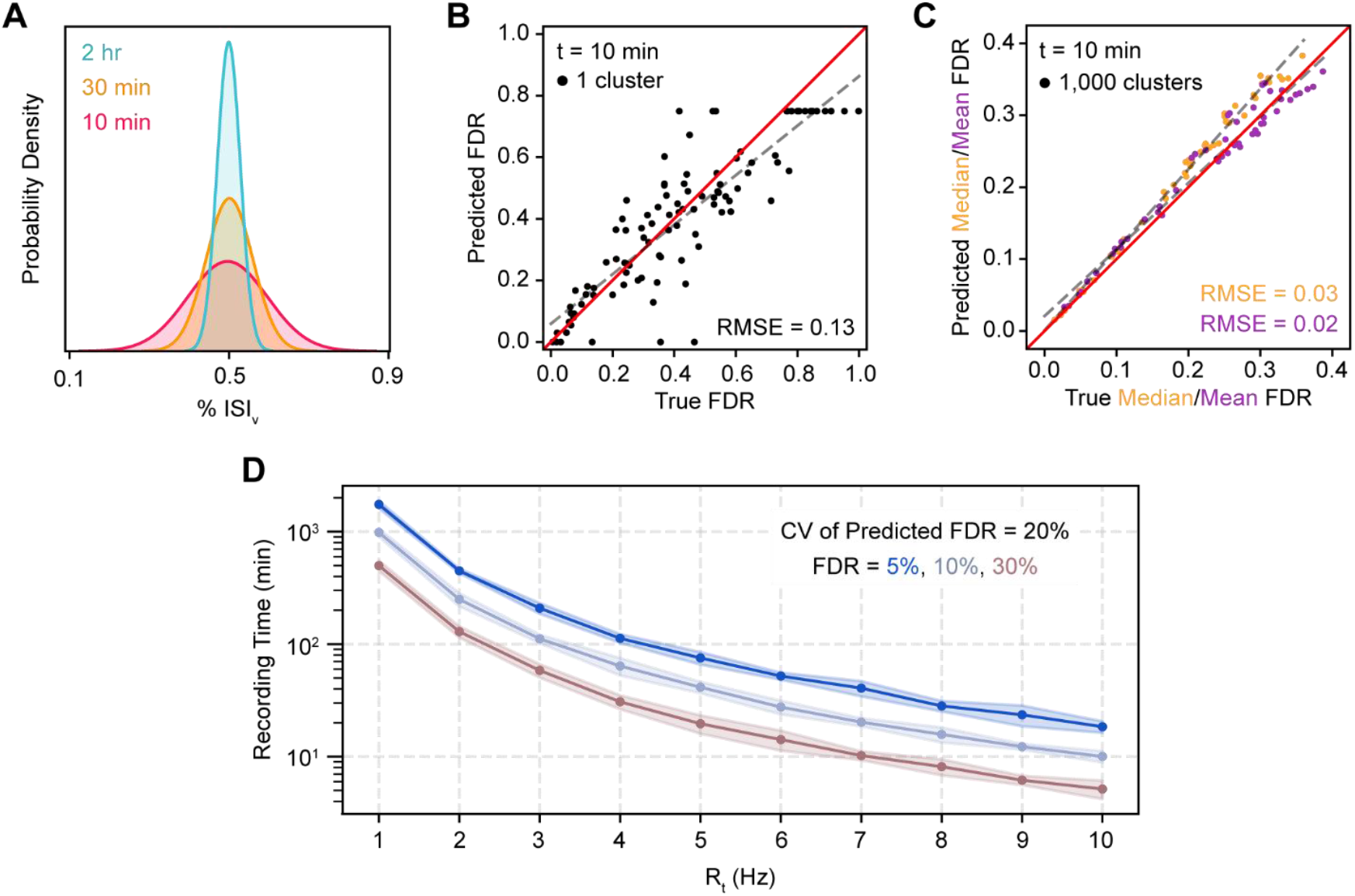
Prediction of FDR for single clusters and populations. (**A**) Probability density functions of observed % ISI_v_ for an example 8 Hz, 15% FDR cluster recorded for varying time lengths. (**B**) Prediction of FDR in single clusters recorded for 10 minutes. All parameters varied simultaneously across a range of physiologically relevant underlying neuronal characteristics (see *Materials and Methods*). Ceiling effect of predicted FDR at 0.75 due to theoretical limit of FDR when predicting with unknown N (see *Materials and Methods*). Red line is the unity line, or perfect concurrence between predicted and true FDR; dashed gray line is the line of best fit. RMSE calculated with respect to the unity line. (**C**) Prediction of median and mean FDR in 1,000 cluster populations. Cluster FDRs are Cauchy-distributed around the population mean. Parameters varied simultaneously and conventions as in (*B*). (**D**) Minimum recording time required for predictions of FDR in a single cluster to have a CV of 20%.

### Calculation of peristimulus time histograms (PSTHs)

PSTHs of cluster responses for *in vivo* electrophysiology data were calculated using bin sizes of 50 ms. Predicted median and mean FDR were found to be largely unaffected by bin size selection in trial-averaged PSTHs. For continuously recorded data, the greatest length of time that could be extracted around each cue without overlapping was used to generate trial-averaged PSTHs.

### Estimation of FDR in publicly available electrophysiology datasets

When predicting FDR using recorded data, only limited information is available about each sorted cluster. 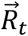and ISI_v_ can be computed directly. τ and 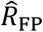, can be estimated, the former through prior knowledge of electrophysiological properties of neurons in the organism and brain region being recorded from, and the latter through examination of other sorted clusters present in the recording session. Estimating N is not straightforward.

To account for uncertainty in these parameters, two approaches can be taken. Appropriate values can be estimated, and simply plugged directly into the equation, or a range of probable values can be input and their results averaged. In this work, for calculation of the values in Table 1, a mixture of both methods was used. An N of both 1 and ∞ were assumed: in the former case, a given sorted cluster was compared to every other cluster in the recording session, while in the latter, it was compared to a single global 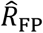 averaged across the session. Clusters for deriving 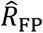 could conceivably be restricted to those on the same or nearby electrode sites, although this was not the approach used here due to uncertainty about each dataset’s probe geometry. A single τ of 2.5 ms was used for every dataset, all of which were composed of recordings from mouse brains. This refractory period was modified by the censor period, when necessary. Censor periods were determined through visual inspection of aggregated per-cluster ISI histograms across the entire dataset. The final FDR is the mean of the (N = 1) cases averaged with the (N = ∞) case. Empirically, we found this equivalent to assuming a single N of approximately 2-3. Our results indicate that FDR estimates are not highly sensitive to the particular choice of N (**Fig. 2C**).

In some datasets, spontaneous activity was recorded, and no cue or event was present to calculate PSTHs around. In such cases, FDR was initially predicted using cluster firing rates calculated across the entire session. However, these predictions were found to be sensitive to bin size used when calculating firing rates, therefore the predicted FDR assuming homogeneous firing is given instead (**Table 1; Eq. 9**).

Due to stochasticity associated with ISI_v_ estimates, observed ISI_v_ values sometimes exceed theoretical bounds given by the chosen parameters, producing imaginary predicted FDRs. In such cases the predicted FDR is capped at its theoretical maximum for a given (assumed) number of contaminating neurons. (**Eq. 1**). This maximum is derived from the fact that given a certain N, if predicted FDR exceeds FDR_max_, a lower FDR could be attained by simply selecting a different neuron within the sorted cluster to be the intended or “true” neuron. When FDR is being calculated by averaging the (N = 1) and (N = ∞) cases, FDR_max_ is the average of these two cases’ maximums: 0.75.

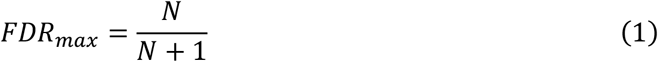

For selecting datasets to examine, only papers published in the last 10 years with publicly available spike-sorted electrophysiology data were considered. No limits or minimums were placed on cluster count, and no sorting methods were specifically included or excluded.

## RESULTS

### The relationship between ISI_v_ and FDR is complex

We sought to understand how the occurrence of ISI violations depends on underlying cluster FDR and how other underlying characteristics of neuronal activity might affect this relationship. To this end, we focused on three variables likely to have a meaningful effect on the occurrence of ISI violations: neuronal firing rate (**Fig. 1A**), temporal correlation of activity amongst the recorded population of neurons (**Fig. 1B**), and the number of contaminant neurons (**Fig. 1C**).

To determine how these variables affect the relationship between ISI_v_ and FDR, we used Monte-Carlo simulations of neural spiking to examine how the relationship between ISI_v_ and FDR might change as a consequence of varying each parameter in isolation. Total cluster firing rate was found to have a dramatic effect on ISI violation production, since overall firing rate played a large role in determining the likelihood that any given FP spike would produce an ISI violation. Critically, clusters with lower firing rates and clusters with higher firing rates could both present with the same ISI_v_, even when they had markedly different underlying FDRs (5% - 50%) (**Fig. 2A**). These results indicated a nonlinear relationship between ISI violation production and firing rate not accounted for by simply dividing the number of ISI violations by total firing rate (ISI_v_). The temporal overlap between TPs and FPs was also found to strongly modulate ISI violation production, although not as strongly as cluster firing rate (**Fig. 2B**). Variable TP-FP covariance (0.8 to -0.5 Hz^2^) altered the probability of any given FP leading to an ISI violation, resulting in three clusters with substantially different FDRs (13% - 36%) presenting with the same ISI_v_ and firing rate. Lastly, number of contaminant neurons was also found to modulate ISI violation production, although not as meaningfully as other variables (**Fig. 2C**). Greater numbers of contaminant neurons increased the odds of ISI violations between pairs of FPs, meaning the same observed ISI violation rate was associated with a slightly lower FDR in cases of multiple contaminant neurons vs. just one contaminant neuron. The dependence of this phenomenon on FP-to-FP violations means its effects only became meaningful at high FDRs (>0.25) and total firing rates (>10 Hz).

### Analytical model of the relationship between ISI_v_ and FDR

We next sought to derive an analytical model describing the dependence of underlying FDR on observed ISI_v_ that incorporates each of these variables. To that end, we first considered a simplified case in which two neurons are each firing homogeneously, or at a constant rate, and spikes from both neurons are being assigned to the same cluster. Each TP spike produces a double sided “violation window” in time. If an FP spike occurs within that window, an ISI violation is observed (**Eq. 2**; **Fig. 1C**).

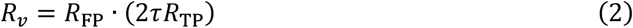

Here, we represent the number of ISI violations observed per second as R_v_, the neuronal absolute refractory period as τ, and the observed firing rates of the two neurons contributing TPs and FPs as R_TP_ (TP rate) and R_FP_ (FP rate). Note that R_TP_ and R_FP_ represent only the rates at which TPs and FPs are assigned to a sorted cluster, not necessarily the total firing rates of the neurons producing the TP and FP spikes. These may be one and the same, e.g., for a neuron contributing TPs with no false negatives. This equation can be solved using the quadratic formula (**Eq. 3**) producing an expression for FDR as a function of ISI_v_, τ, and R_t_ – the total observed firing rate of the cluster – using a few simple relationships (**Eq. 4-6**). The larger root is ignored, i.e. the term under the square root is subtracted and not added, because the neuron with the most spikes in the cluster is *de facto* considered the TP-contributing neuron (see **Materials and Methods**).

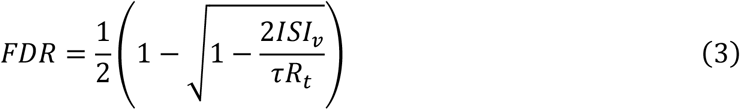

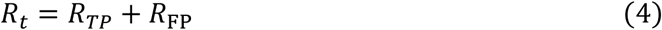

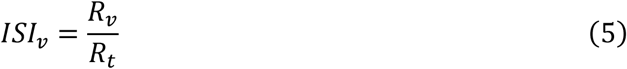

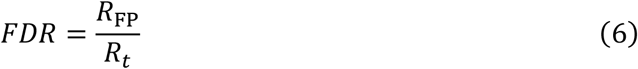

Some spike sorting algorithms make use of a censor period, whereby spikes detected within a certain minimum distance, τ_c_, of another spike are ignored. In such cases, the size of the violation window produced by each true positive spike is shortened by this censor period producing a new effective refractory period: τ_e_ = τ – τ_c_. Implementation of τ_e_ produces (**Eq. 7)**, which is equivalent to a rearranged form of the equation derived in (Hill et al., 2011).

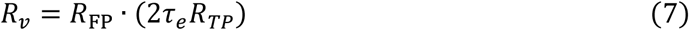

At the other extreme, consider a situation where the spikes comprising a cluster are generated by an infinite number of neurons. In this case, FPs are capable of producing ISI violations with other FPs, necessitating the addition of a second term wherein FP spikes now produce double-sided violation windows as well. This term is scaled by a factor of one half to prevent double counting of FPs producing ISI violations with one another. Implementation of this term produces (**Eq. 8**), which is equivalent to a rearranged form of the equation derived in (Llobet et al., 2022). This equation can be solved for FDR using the quadratic formula as previously.

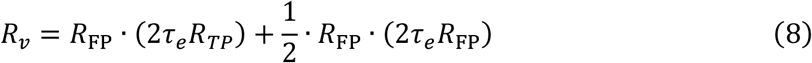

For an unspecified number of contaminant neurons N, an additional scaling factor of (N – 1)/N can be added to the FP-FP ISI violations term (**Eq. 9**). This factor can be interpreted as the fraction of all FPs available for any given contaminant neuron’s spikes to produce ISI violations with, e.g., 1/2 for N = 1, 2/3 for N = 2. This equation, like previous iterations, can be rearranged to solve for FDR with an additional dependence added on N.

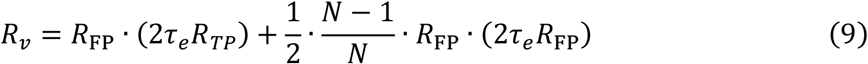

In the more biologically relevant case of inhomogeneous firing, or neural spiking rates that vary over time, R_TP_, R_FP_, R_t_, and R_v_ can all be considered not as constants but as functions of time (i.e. vector quantities). While this spiking nonstationarity must be taken into account, a time-varying estimate of FDR would be a needless level of granularity and also highly inaccurate given the stochastic nature of neuronal spiking and ISI violations, so of primary interest is a time-averaged estimate of the relationship between violation rate 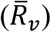 and underlying variables. In this case, the rate of violations depends not on the product of the average values of R_TP_ and R_FP_, but on the expected value of their element-wise product, 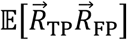:

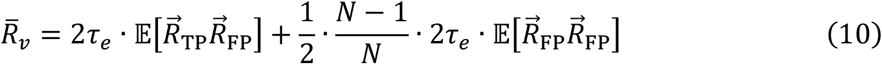

For two vectors representing firing rate over time 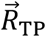 and 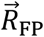of length n elements, this expected value can be calculated as follows:

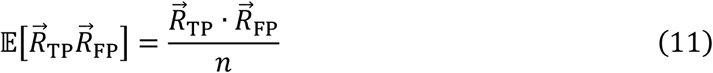

Equation 10 can then be solved for 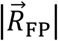, the vector magnitude of 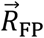, using the quadratic formula (**Eq. 12**). Unit vector 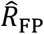 can subsequently be scaled by 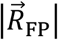, averaged, and finally divided by the mean total firing rate of the cluster to obtain a time-averaged estimate of FDR (**Eq. 13**).

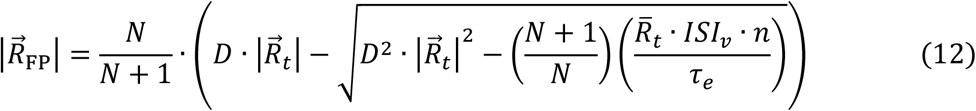

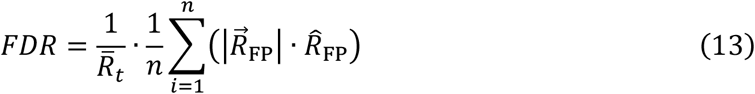

Here, D corresponds to the dot product of the unit vectors representing total cluster spike rate and cluster FP spike rate, 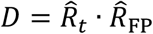. This can be thought of more generally as representing the degree to which the time-varying total cluster spike rate temporally overlaps with the time-varying FP spike rate. This final equation depends upon a number of parameters specific to each cluster to obtain a single FDR estimate: (1) the effective refractory period τ_e_, (2) the temporal distributions of activity in the cluster of interest 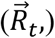 and (3) of other clusters contributing FP spikes 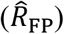, (4) the observed ISI violation fraction ISI_v_, (5) the number of contaminant neurons N. For information on how one can estimate these parameters from experimental data, see **Materials and Methods**.

### Prediction of FDR in silico

To both assess our model’s predictive power as well as more generally illustrate relationships between model variables, we next simulated neural spike trains while varying all relevant parameters across a range of biologically relevant values and then attempted to predict FDR from the observed ISI_v_. In this case, parameters like N and 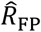 that may normally have associated uncertainty are known exactly. Simulating long periods of time (up to 28 hours of recording time) also enables highly accurate estimates of ISI_v._

For both homogeneous and inhomogeneous firing we found that analytical FDR predictions closely approximated the true underlying FDRs (**Fig. 3**). For homogeneous firing specifically, observed ISI_v_ was found to strongly depend on FDR and total cluster firing rate, as expected (**Fig. 3A**). A linear dependence was observed of ISI_v_ on firing rate at fixed FDR and contaminant neuron counts, despite ISI_v_ often being assumed to already have normalized for cluster firing rate. Furthermore, FDR was found to scale quadratically with increasing ISI_v_ at a fixed firing rate and contaminant neuron count (**Fig. 3B**). ISI_v_ values of 0.1-1% represent typical thresholds in literature for considering a cluster well isolated (Boucher et al., 2023; Chandrasekaran et al., 2017; Jadhav et al., 2009; Roy & Wang, 2012; Wright et al., 2021; Zhao et al., 2023). Yet, our results indicate that FDRs associated with these ISI_v_ values vary considerably with the firing rate of the cluster in question (**Fig. 3A-B**). As an illustration, an ISI_v_ of 0.5% reflected a desirable 5% FDR for a cluster firing at 20 Hz, or a much higher 50% FDR for one firing at 3 Hz. In general, contaminant neuron count was of limited consequence unless the cluster in question had both a high firing rate and high FDR, making FP-FP violations frequent enough to meaningfully affect overall ISI violation incidence.

For clusters that fired inhomogeneously in time, errors in FDR predicted with the homogeneous equation (**Eq. 9**) scaled linearly with the temporal covariance of TPs and FPs (**Fig. 3C**). Positive covariance increased the ISI violation rate at the same FDR, resulting in overestimation of FDR, while negative covariance conversely decreased ISI violations, resulting in underestimation of FDR. When TP and FP spike rates do not covary in time, the relationship between ISI_v_ and FDR mimics the homogeneous case even if the generation of TPs and FPs individually may not necessarily be homogeneous. Only under these conditions did the homogeneous and inhomogeneous predictions agree. When inhomogeneous firing was appropriately taken into account (**Eq. 10**), predicted FDR closely approximated true FDR regardless of the covariances in neuronal firing.

When FDR was predicted from ISI_v_ using **Eq. 10**, predicted FDR maintained high agreement with true FDR across the entire space of parameters investigated (**Fig. 3D**; RMSE = 0.02).

### Prediction of FDR under realistic conditions

Our model performed well when ISI_v_ and other parameters were known exactly, although benchmarking simulations indicated that predictions of FDR are sensitive to small changes in ISI_v_ (**Fig. 3**), particularly at lower firing rates (**Fig. 3B**). We next wanted to determine whether FDR could be accurately predicted from recordings of finite duration with noisy ISI_v_ estimates and when exact values of N and 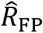 are unknown. Simulating spiking for finite durations, we found that observed ISI_v_ values were normally distributed around their true values (**Fig. 4A**), with increasing recording time decreasing the variance of this distribution, as expected.

To assess the effect of noisy ISI_v_ estimates on FDR predictions, we again simulated spiking while simultaneously varying all previously described parameters, but this time recording duration was restricted to 10 minutes (**Fig. 4B-C**). In this case, we found that FDR predictions for individual clusters were substantially less accurate (RMSE = 0.13), although they did not deviate systematically from true FDRs (**Fig. 4B**). Even with this highly restricted recording time, FDR population statistics could be estimated across a set of clusters with high fidelity so long as a sufficient number of clusters were sampled (**Fig. 4C**). For example, median FDR and mean FDR could be predicted with RMSEs of 0.03 and 0.02 respectively across 1,000 clusters. Predicted median FDRs were slightly overestimated at high true median FDR (>0.25).

We next sought to determine the duration of neural recordings necessary to obtain accurate estimates of ISI_v_, and thus, FDR, in single clusters. To accomplish this, we simulated clusters with various firing rates and FDRs, and determined the recording time sufficient to produce a coefficient of variation (CV) of the predicted FDR of 20% (**Fig. 4D**; e.g. 50% ± 10% or 5% ± 1%). Surprisingly, we found that clusters with firing rates of 1-2 Hz required observations across hundreds of minutes to produce accurate estimates of single cluster FDR, an infeasible recording time in many common experimental paradigms. For clusters with firing rates greater than 5 Hz, FDRs between 5-30% could be estimated accurately using tens of minutes of spiking data.

We used our model to estimate the FDRs of clusters contained within 12 publicly available datasets that included spike-sorted electrophysiology recordings (**Table 1**). Given limitations associated with single cluster FDR predictions (**Fig. 4B**), estimated median and mean FDR were reported across all clusters present in each dataset. An average median FDR of 12.9% ± 13.5% (s.d.) was observed along with an average mean FDR of 24.1% ± 9.2% (s.d.). No obvious correlations between cluster count, spike sorting methodology, or recording technology and dataset FDR were observed. Median estimated FDRs were consistently lower than mean estimated FDRs. This implies that cluster FDR distributions in recorded electrophysiology datasets tend to be right-skewed, composed of a large proportion of clusters possessing FDRs closer to 0 as well as a broadly distributed complement of more contaminated clusters, some of which potentially reaching FDRs well above 0.5.

Importantly, no attempts were made to curate clusters included in analysis from each dataset; all available sessions and clusters were assessed, with the exception of clusters labeled “multi” or “MUA” in datasets with cluster quality annotations. It’s possible that some datasets were precurated, with clusters discarded according to some exclusion criteria prior to being uploaded, while others were shared un-curated. Depending on the nature of the scientific question being addressed, better cluster isolation is undoubtedly a larger priority for some datasets than others. For these reasons and due to uncertainty in assessing censor period in each dataset, we emphasize that these findings should serve generally as an overall survey of the range of expected contamination levels in datasets produced using widely used methods rather than as a commentary on individual datasets or spike sorting methodologies.

## DISCUSSION

Inter-spike interval violations are the most commonly employed metric of accuracy in spike sorting, serving as an indication of the false discovery rate (FDR) – the rate at which spikes are erroneously assigned to the wrong cluster. Here, we used Monte Carlo simulations to demonstrate that the inter-spike interval violation rate (ISI_v_) is related to FDR through a complex relationship that depends on many factors, including (1) the neuronal firing rate, (2) the temporal correlation in activity between neurons contributing to a cluster, and (3) the number of neurons contributing spikes to a cluster - in that order of descending importance. We derived an analytical model that can be used to predict FDR from ISI_v_ that incorporates these factors and determine the accuracy with which FDR can be inferred during finite-length recordings at the level of single clusters and datasets. Finally, we used this model to assess the FDR of clusters contained in publicly available spike-sorted electrophysiology datasets to provide bounds on the accuracy that can be reasonably expected by experimenters. Our study makes four central contributions.

First, we derive an analytical model that can be used to estimate FDR from ISI_v_ accurately across a broad parameter space.

Second, we explicitly demonstrate that FDR is not linearly related to ISI_v_, but depends critically on the total cluster spike rate. While this dependence can be inferred from previous work (Hill et al., 2011; Llobet et al., 2022), our results underscores the inappropriateness of using ISI_v_ as an inclusion criterion for single clusters – which is still a common practice in many studies using spike sorted data. Across a common range of firing rates (∼2 to 12 Hz), clusters with the same ISI_v_ can be associated with both low (∼5%) and very high (∼50%) FDRs (**Figs. 2A, 3A**).

Third, we find that estimates of FDR at the single cluster level are noisy due to the stochasticity of ISI violations as well as uncertainty in cluster-specific parameters (**Fig. 4B**) – but estimates at the population level are highly robust (**Fig. 4C**). As a point of reference, our results suggest that ISI_v_ can be estimated accurately enough to predict single-cluster FDRs within 20% of their true values in one-hour recordings only when firing rates are greater than ∼5 Hz. Even for clusters that meet these requirements, however, experimental uncertainty in parameters like FP-TP covariance and contaminant neuron counts make single cluster FDRs difficult to predict with high confidence. Alternatively, population-level statistics of FDR, obtained by averaging across all the clusters in a dataset, can be accurately predicted with recording durations as low as 10 minutes.

Finally, we predict FDR on the basis of ISI violations in publicly available datasets for the first time. FDR population statistics covered a wide range (median: 3.1 – 50.0%; mean: 12.5 - 44.3%) and could be estimated with low standard error (s.e. median: 0.2-1.3%; s.e. mean: 0.2-1.0%). Datasets with low FDR were not associated with any obvious external features of data collection, spike sorting methodology, or recording technology and the variance in FDRs across datasets is likely a function of the variable importance of low FDR for the scientific goals of individual studies.

### Monte Carlo simulations for validating prediction of FDR

Given the absence of high cluster count spike-sorted extracellular neural recordings with associated per-neuron ground truth patch clamp data, Monte Carlo simulation presents itself as an attractive tool for studying the theoretical mechanisms by which a given level of contamination in a sorted cluster translates into observed ISI violations. The fundamental assumption of all simulations in this work is that neural spiking resembles a collection of independent Poisson processes, a common assumption that has been validated across a number of organisms, brain regions, and behavioral contexts (Abbott & Dayan, 2005; Roxin et al., 2008; Shinomoto & Tsubo, 2001; Tolhurst et al., 1981; Werner & Mountcastle, 1965). Some work has, however, pointed to the possibility of non-Poisson firing in certain brain regions (Maimon & Assad, 2009; Swindale et al., 2023); care should be taken in application of this work to data obtained from these areas. Beyond the foundation of independent Poisson firing, only aspects of neuronal firing and spike sorting thought to potentially be relevant to ISI violation production were modeled, namely varying firing rate amplitudes, temporally inhomogeneous firing, and differing contaminant neuron counts. If an additional aspect not accounted for in this description plays a significant role in determining the relationship between ISI_v_ and FDR, then *in silico* validation may not reflect true congruence between analytical prediction and reality.

### Best practices for using ISI violations in spike sorting

When attempting to determine the success of spike-sorting operations *post-hoc*, ISI violation fraction is frequently used as a per-cluster inclusion criterion. However, unless a cluster is recorded for a long enough time period given its firing rate and true FDR and difficult-to-estimate cluster-specific parameters are known, it can be difficult to predict FDR using ISI violations at the single cluster level with high confidence. Use of ISI violation fractions in this way can easily result in situations where highly contaminated clusters are erroneously kept while less contaminated clusters are discarded. We posit that the most straightforward and robust use case for ISI_v_ is as a tool for predicting population-level statistics of FDR when coupled with a sound theoretical understanding of how cluster contamination translates into ISI violations (**Eq. 10, 12, 13**). Investigators can obtain an accurate estimate of median and mean cluster FDR across a session or dataset and then decide whether they are satisfied with these levels of cross-contamination, or if additional effort to improve cluster isolation is needed.

When curating spike-sorted data, it is critical that both algorithmic and manual sorters do not specifically remove individual spikes that generate ISI violations. Typically, only a small fraction of contaminant spikes produce ISI violations; targeted removal of spikes producing ISI violations can reduce ISI_v_ substantially without meaningfully reducing the FDR, thus producing clusters that seem well isolated based on their ISI_v_, even when they are not. In effect, this practice does not accomplish anything except eliminating the predictive power of ISI_v_ for underlying FDR.

### Current state of spike sorting predicted using ISI violations

This work estimates an average median FDR of ∼13% and an average mean FDR of ∼24% in publicly available electrophysiology datasets. The lower median FDR compared to mean FDR across virtually all datasets examined indicates right-skewed FDR distributions. This likely arises as a consequence of FDR having a theoretical floor of 0, with most datasets having many cluster FDRs close to this floor. It may also be a consequence of most spike sorting datasets being composed of two distinct types of clusters: relatively easier to sort clusters whose FDRs typically fall close to 0 and relatively harder to sort clusters whose FDRs are likely to fall more broadly over the theoretical range of FDR (0-1).

In the absence of any clear rationale for variance in FDR as predicted using ISI_v_ among publicly available spike-sorted datasets (**Table 1**), degree and execution of manual curation presents itself as a promising explanatory candidate. While modern spike-sorting algorithms serve as an excellent basis for sorting vast quantities of electrophysiology data, many investigators still manually merge, split, and discard algorithmically obtained output clusters to further improve cluster isolation. Time, effort, and skill applied to manual curation are difficult to quantify and therefore unlikely to be reported in literature, although such differences are likely to have a material effect on the final quality of cluster isolation. The median and mean FDRs of the datasets examined here as well as the tendency toward right-skewed FDR distributions support the idea that all datasets are composed of both well and poorly isolated clusters.

### Necessity of high-quality spike sorting

It has been posited that well-sorted clusters are not a necessity for many types of neural data analyses, particularly those concerned with studying population dynamics (Christie et al., 2014; Trautmann et al., 2019). In some applications, however, well-isolated clusters remain a critical precondition for answering relevant neuroscientific questions. Characterizing the responsivity of specific cell types that have been identified on the basis of genetic expression, projection target, waveform shape, or activity *in vivo* represents an expansive line of inquiry wherein high-quality cluster isolation is key (Deubner et al., 2019; Ding et al., 2022; Estebanez et al., 2017; E. K. Lee et al., 2021; Takatoh et al., 2022). Ultimately, the level of cluster isolation necessary for a given study is highly dependent upon the biological questions of interest. The work herein aims to clarify the relationship between ISI violations and cluster contamination, as well as provide a tool by which overall spike sorting quality can be quickly assessed with a direct, interpretable, and accurate metric, thereby streamlining assessments of sorting performance and increasing confidence that desired cluster isolation levels have been reached.

## ACKNOWLEDGEMENTS

We thank Munib Hasnain, Maria Moya, Yujin Han, Jackie Birnbaum, and Will Cunningham for their helpful comments on the manuscript. This work was supported by 1R01NS121409.

## AUTHOR CONTRIBUTIONS

J.P.V.: Designed and performed research, analyzed data, wrote and edited the manuscript M.N.E.: Designed research, wrote and edited the manuscript

